# A dispersal model for *Cenchrus setaceus* (Poaceae) in Mexican urban context

**DOI:** 10.64898/2026.10.17.759854

**Authors:** Rosalio Reyes, Vincenzo Bertolini, Rafael Barrio-Paredes, Jorge José Ochoa-Bonilla, Tomás Martínez-Trinidad

## Abstract

Urban environments suffer from multiple ecological issues, such as the invasion of alien. In Mexico, *Cenchrus setaceus* (African fountain grass) has been introduced as an ornamental species in the 1990s spreading all around the country. The aim of the present study is to propose an invasion models, based on the temporal dispersion of *C. setaceus* seeds using field data from the metropolitan area of Guadalajara, Jalisco, distinguished between planted individuals from self-propagated ones, quantifying seeds fertility *ex situ*. Model indicate that the current infested area of the Guadalajara metropolitan area could reach 51-89% within 100 years. Applied to three cities in Chiapas, the model shows limited spread over 50-year period from existing *foci* but strong sensitivity to new ornamental introductions. An analysis of containment strategies estimates an invasion front velocity of 0.13 km/year and identifies a 3-km-wide buffer zone as necessary to halt spread over a 25-year horizon.

## 1 Introduction

Cities are facing growing socio-environmental challenges arising from implementation and spreading of global urbanization processes [1]; indeed, it is estimated that by 2050, 78% of the global population will reside in urban areas [2]. Urban green areas—including trees and vegetated public spaces—are fundamental socio-physical elements for urban sustainability, simultaneously contributing to human well-being, environmental regulation and quality of life [3, 4]. These urban areas are often plagued by serious environmental issues related to the spread of alien species, frequently introduced for esthetic reasons, which are capable of altering ecological balances, sometimes irreversibly, even beyond city boundaries, as documented in the case of African fountain grass (*Cenchrus setaceus* (Forssk.) Morrone) in various regions of the world [5]. This species is an exotic herbaceous plant of the Poaceae family, native to Northern Africa, East Africa and the Arabian Peninsula [6], commonly used as an ornamental plant or forage in various parts of the world. This plant turns out to be a very aggressive colonizer almost everywhere, including Mexico [7, 8]. Today, it is considered cosmopolitan, distributed across all continents. The species flourishes within an extensive climatic range, as long as the annual precipitation does not exceed 1,270 mm and the temperature does not fall below 0 ^*°*^C [9]. It is a perennial plant that can live for up to 20 years and produces numerous seeds, which could remain viable in the soil potentially for up to 6 years or longer [10]. It is also highly resilient to the impact of the fire [11]. Since it adapts to degraded and poor soils and drought conditions, it has become invasive in several countries degrading both the soil and biological diversity as well as displacing native grass species [12]. Its presence and negative impacts have been reported in Hawaii [13, 14], the mediterranean region [9, 15] and Mexico [5]. Urban environment are a relevant as spreading centers of exotic species because of ornamental uses [16]. Human population density appeared as the strongest predictor of the occurrence and richness of naturalized species, highlighting the role of human activity in promoting invasions in the american continent [17]. The urban habitat simulates the pre-desert areas of the Sahara where this species come from, in urban heat islands (UHIs) and in areas with continuous soil disturbance from infrastructure development, significantly accelerating the invasion [18]. Notice that roads facilitate the actual propagation of seeds because the passage of vehicles creates air currents that can easily transport the tiny seeds, while the degraded soil at the edges of the roads is easily colonized by the species [9]. All these characteristics make the species an organism capable of modifying the ecology of the colonized territory [19].

On the other hand, We should point out that one can monitor a biological invasion in real time with a citizen-science platform such as iNaturalist, that allows us to identify, record and share observations of biological organisms, providing georeferenced data of species occurrences. The first iNaturalist recorded sighting in Mexico for C. setaceus was in 2012 in Ciudad Victoria, Tamaulipas [20]; as of January 1st, 2016, a total of 12 points of interest has been recorded in Mexico, making the total of 2,152 records up to date [20]. Particularly, in the Guadalajara metropolitan area in Jalisco (Figure 1b), the first records of *C. setaceus*, reported on the platform in 2019, identified 4 points of interest [20]. According to local reporters, *C. setaceus* was introduced to Guadalajara in early 1996 as an ornamental plant for the urban landscape, with the aim of redesigning the cityscape for the sporting event “*Universiada Nacional* “, whose final stage tooks place in this city [21, 22].

**Figure 1.**
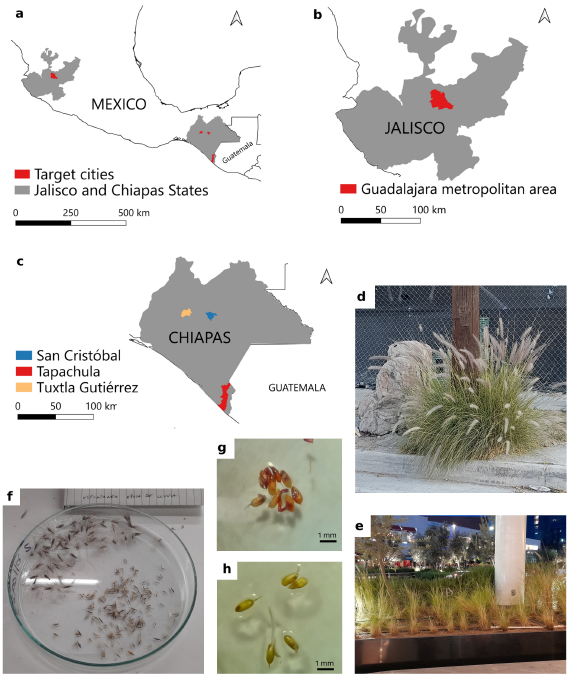
Geographical localization, examples of invasion, seed counting and fertility. a. The entire Mexican territory, highlighting the states of Jalisco and Chiapas. b. Detail of the state of Jalisco: Guadalajara metropolitan area (modeling site). c. Detail of the state of Chiapas: San Cristóbal de las Casas, Tapachula, and Tuxtla Gutiérrez (cities subject to projection). d. Self-propagated individual. e. Planted individuals. f. Spikelets and seed counting. g. Tetrazolium test. h. Germination test

In general, classical mathematical models based on reaction-diffusion equations predict that an invasion advances as a traveling wave front at a constant asymptotic speed [23, 24]. However, propagation of invasive plant shows a spotted, patch pattern, driven by long-tailed dispersal distributions [25, 26], which have been described by models based on diffusion kernels or cellular automata that evolve through probabilistic rules and have recently been calibrated against real data to project medium and long-term spread [27]. For *C. setaceus*, correlative species distribution models (SDM) have been developed for the Canary Islands [28] and Mexico [5], and a nested hierarchical SDM framework, combining global and regional climatic niches, has been developed [29]. These models delimit climatically suitable areas for the weed and describe its potential distribution throughout the territory, but do not describe the temporal evolution of spread from a localized source. With the aim of modeling the invasion dynamic in Mexico, particularly in the Chiapas state (Figure 1c), we propose a dynamical model for *C. setaceus*, based on the spread observed in recent decades in the city of Guadalajara, Jalisco, Mexico (Figure 2a), where we georeference the infestation *foci*. For each *focus*, we recorded the number of individuals and the mean number of spikes per individual. We also distinguish between two types: planted by humans and self-propagated *foci* (Figure 1d, e). We estimate seed production and its viability. We then develope a model consisting of two modules. The first is an exponential dispersal-kernel module and the second is a local stochastic establishment module. We calibrate the model using the data collected in Guadalajara and projected the infestation scenario over 50 and 100-year horizons for the city. We also geo-referenced the presence data of *C. setaceus* in three Chiapas cities, where the infestation is currently beginning. We projected two scenarios: the current initial conditions that remain unchanged over 50 years and the continued planting of the species as an or-namental plant. We project the scenario over a 200-year horizon in Mexico using iNaturalist data. Finally, we analyze strategies to repel or contain the weed.

**Figure 2.**
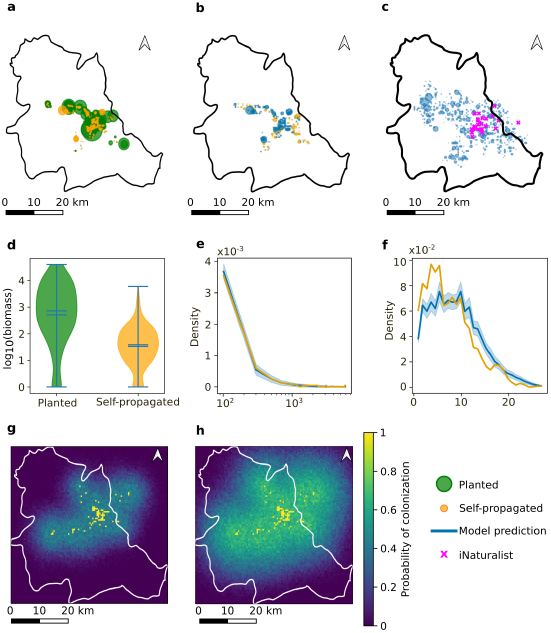
Data collected in Guadalajara and model projections. a. Scatter plot of the collected data in Guadalajara. Green indicates planted and orange indicates self-propagated data. The size of the circle is proportional to the biomass size. b. Comparison of real data and simulated results. In orange, self-propagated data recorded in Guadalajara. In blue, one run of the model by 26 years. We show the points inside concave hull of the complete real dataset. c. If we plot all the points predicted by the model, we can observe that the data expand beyond of sampled area by we. In magenta, we observe presence data obtained from iNaturalist. d. Distribution of the collected biomass. Notice the logarithmic scale in the y axis. e. Comparison of density distributions functions of biomass for the self-propagated and model prediction cases. f. Comparison of density distributions of the distance between self-propagated *foci* for both cases. Projections for Guadalajara in 50 and 100 years. Probability of pixel colonization after 50 (a) and 100 (b) years. Yellow pixels indicate the source.

## 2 Results

### 2.1 Estimation of seed productivity, fertility and germination rate

During the dry season, self-propagated plants showed an average of 182 spikelets per spike (std.dev. = 52.5), and an average of 54.5 seeds per spike (std.dev. = 24.4). For planted plants, an average of 157 spikelets per spike (std.dev. = 51.6) and an average of 45.4 seeds per spike (std.dev. = 23.5) were recorded. During the rainy season, we obtained the following results: for self-propagated plants, an average of 173 spikelets (std.dev. = 35.1) and an average of 70.3 seeds per spike (std.dev. = 30.1). For planted plants, we obtained an average of 159.1 spikelets (std.dev. = 19.4) and an average of 35.4 seeds per spike (std.dev. = 17.9). The total number of spikes recorded in the census was 476,142 for planted plants and 37,840 for self-propagated ones. Under a conservative annual reproductive cycle with 500,000 spikes per year (assuming a single annual cycle) and a conservative average of 50 seeds per spike, yielding a production of 25,000,000 seeds per year.

The tetrazolium test showed that seeds collected during the dry season reached 87.7% viability after 24 hours from planted individuals and 86.8% from self-propagated ones. During the rainy season, the same data showed values of 83.6% and 88.7%, respectively. Germination tests conducted 48 hours after sowing seeds, collected during the dry season showed a germination rate of 78% for planted and 75% for self-propagated individuals. During the rainy season, the same figures are 58.3% and 43.3% for planted and self-propagated, respectively.

Under a baseline scenario, assuming a 50% germination rate for the seeds, we would have 12,500,000 new seedlings per year (always assuming, by default, only one seed production per year). According to an analysis conducted in Europe, germination is a key factor in the naturalization of exotic species [30]. The physical distance between occurrences of planted and self-propagated averages 6.13 km, with a standard deviation of 4.84 km, a minimum distance of 0.007 km and a maximum distance of 28.45 km.

### 2.2 Data are described by a lognormal distribution

We analyzed candidate probability distributions for our data. Evaluations were conducted separately on the complete dataset as well as on the individual sub-datasets (planted and self-propagated) using multiple criteria provided by the `fitter`function within the Fitter Python package.

Across all evaluated datasets, a consistent pattern emerged. Bounded distributions, such as the Beta distribution, yielded the lowest Sum of Squared Errors (SSE) over the central mass of the empirical distributions, whereas the Lognormal distribution consistently yielded the lowest Akaike Information Criterion (AIC) and Bayesian Information Criterion (BIC) values.

Based on these information criteria, the Lognormal distribution was selected for the simulation framework. We assume that this distribution function corresponded to 26 time steps because we assume that the planted source was established 26 years ago.

Let *B* denote the biomass variable (see eq. 8). We assume that

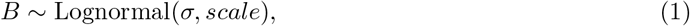

which is equivalent to

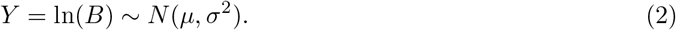

The scale parameter (eq. 1) used by the `lognorm` function in `scipy.stats` is related to the normal-distribution parameters by *scale* = *e*^*µ*^, while the shape parameter s corresponds to *σ*. In the general parameterization of `scipy.stats.lognorm`, the `loc` parameter introduces a location shift according to

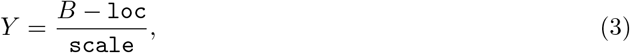

where Y follows a standardized lognormal distribution [31]. In our simulations, `loc=0`

### 2.3 Model fitted to Guadalajara data

We assumed that the annual biomass increment (*B* in Equation (8)) follows a lognormal distribution. We used *σ* and *scale* as optimization parameters. We also used the spatial decay parameter *α* from the exponential kernel (Equation (5)), together with the probability thresholds *θ*_1_ and *θ*_2_, as optimization parameters using the objective function in Equation (11). Starting from the planted data, we ran the simulation for 26 time steps and obtained the following optimized parameter values:

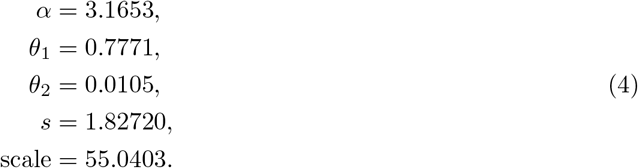

We fixed *loc* = 0. The parameters suggest that once seed arrives, the probability of reaching adulthood is approximately 10%.

Using the parameters from Eq. 4, we simulated weed propagation over a period of 26 years. Figure 2b shows that the resulting distribution exhibits a spotted pattern (blue dots) which closely resembles the observed data (orange dots). Because the field survey was not exhaustive—that is, the large number of locations recorded as absences, reflects unsampled areas rather than confirmed absence of the weed—we calibrated the model only within the concave hull (see Methods), yielding the pattern shown in Figure 2b. Nevertheless, the model predicts the presence of the weed beyond the concave hull, as shown in Figure 2c. In particular, it predicts the occurrence of the weed outside the city boundaries. By analyzing iNaturalist records, we found observations of the weed in several regions where the model predicted its presence but where no field surveys had been conducted. These independent iNaturalist records therefore support predictions of the model.

To evaluate whether the model reproduced the observed distribution of local biomass intensity, we compared the empirical distribution of biomass per pixel from the self-propagated dataset with the distribution generated by the stochastic model. We performed 100 independent model realizations using different random seeds. For each realization, pixel intensities within the concave hull were extracted and used to compute a normalized histogram. The model distribution was estimated by averaging the normalized histograms across all realizations, accounting for stochastic variability in the simulation outcomes. In Figure 2e the model distribution closely resembles the empirical pixel intensity of self-propagated data, showing the invasion in the urban fraction of the metropolitan area of Guadalajara.

We also evaluated whether the model reproduced the spatial organization of colonized areas by comparing the distribution of pairwise distances between occupied pixels in the observed and simulated landscapes. Occupied pixels were defined as cells with positive values within the concave hull. For the observed self-propagated dataset, all unique pairwise distances between occupied pixels were calculated in kilometers. For the stochastic model, the same procedure was applied to 100 independent realizations, and the resulting normalized distance distributions were averaged to account for stochastic variability. The model reproduced the overall spatial organization of colonized cells, with observed and simulated distributions showing similar patterns (Figure 2f). Both distributions were characterized by a high frequency of short-range distances and a gradual decay toward larger spatial separations. However, the observed dataset showed higher densities at short distances, indicating a stronger local aggregation than that produced by the model. Despite this difference, the model captured the general spatial aggregation structure of *C. setaceus* populations.

Once the model was validated, we used it to project future colonization scenarios. We evaluated the probability of colonization after 50 and 100 years after the initial record in 1996 (Figure 2g, h). For each scenario, weed expansion was simulated using the planted matrix as the initial condition for 50 and 100 time steps. Each scenario was independently simulated 100 times, and the colonization probability of each pixel was estimated as the proportion of simulations in which the pixel was occupied.

The current observed distribution of the grass, combining planted and self-propagated occurrences, covers approximately 2.3% of the metropolitan area. After 50 years, the model predicts a spatially heterogeneous expansion pattern. When considering pixels with a colonization probability greater than 30%, the predicted occupied area increases to 32% of the city area. However, under a more conservative threshold of 50% colonization probability, only 1.42% of the city area is consistently colonized across stochastic realizations. This difference between thresholds indicates that a large proportion of the predicted expansion corresponds to areas where colonization is possible but strongly influenced by stochastic dispersal events.

After 100 years, the predicted occupied area increases substantially, reaching 89% of the city area when considering a 30% colonization probability threshold and 51% when considering a 50% threshold, indicating a progressive increase in the spatial extent and certainty of weed expansion over time.

### 2.4 San Cristobal, Tapachula and Tuxtla Gutiérrez projections

The presence data recorded from San Cristobal de las Casas, Tapachula and Tuxtla Gutiérrez in Chiapas were used to project the expansion of the weed in areas where the presence is lower than in Guadalajara. In these cities, we recorded positions without recording biomass size. Therefore, we assigned sizes through random sampling of the Guadalajara data. We then used our model to analyze the scenarios over 50 years.

In San Cristóbal, we recorded six points, and the model predicts that 1% of the entire municipality area will be infected in 50 years (Figure 3a). In Tapachula, we recorded only one point of weed presence. The introduction of this alien species is in an incipient stage. The model predicts an infestation area of 0.1% of the entire municipality area, with a probability of 0.5, 50 years later (Figure 3b). In Tuxtla Gutiérrez, three points were recorded. The model predicts an invasion of 0.2% of the entire municipality area (Figure 3c). It is important to remark that a large proportion of these municipalities has an agricultural land vocation while the model shows the invasion entirely in the urban fraction.

**Figure 3.**
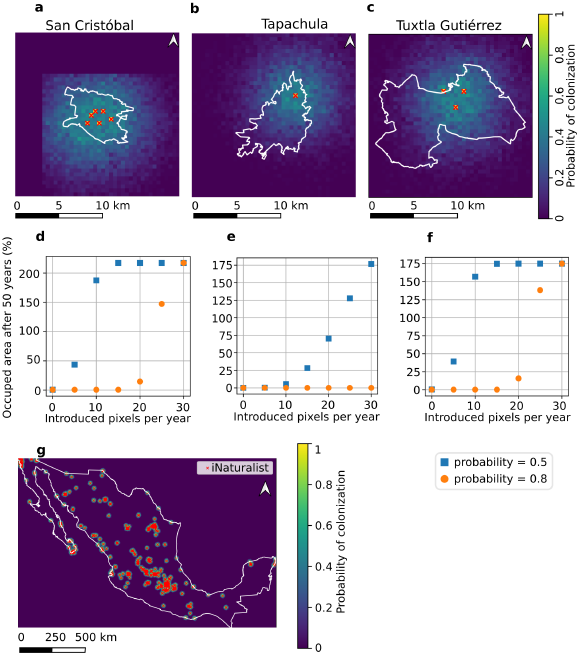
Projections in three cities of Chiapas and Mexico. Probability of pixel colonization in municipality of San Cristóbal de las Casas (a), in municipality of Tapachula (b), and in municipality of Tuxtla Gutiérrz (c). Red x represents initial source. Projections of occuped area after 50 years if we consider annual introductions of new infestation *foci*. d. in municipality of Tapachula, e. in municipality of Tuxtla Gutiérrez, f. in municipality of San Cristóbal de las Casas. Occupied area with a probability of 0.8 (blue) and 0.5 (orange). g. Projection over 200 years in the Mexican territory. Red x indicates the sources from iNaturalist database. After 200 years, the infestation will occupy approximately the 11% of the Mexican territory.

The projections are based on the supposition that expasion is due to the initial infestation *foci*. However, *C. setaceus* is widely used as ornamental plant, and human activity can establish new infestation *foci* that we do not consider in our simulations. In order to take into account these new *foci*, we simulate *n* new introductions per year, with *n* ∈ [5, 10, 15, 20, 25, 30] (Figure 3d, e, f). New introductions were considered in different pixels. The size of the new introductions represents 1% of the biomass size observed in Guadalajara. We observed that in San Cristóbal and Tuxtla, simulation with 25 new infestation *foci* per year predict an infested area of 50% of the cities in 50 years with a probability of 0.8. However, 5 new infestation *foci* generate an infestation of 50% of the total area with a probability of 0.5 in the same period. The case of Tapachula shows that a large initial condition is more important than small infestation points, since the scenario after 50 years is determined by the initial condition rather than by the new small infestation points.

### 2.5 Projections for Mexican territory

The iNaturalist database records the presence of *C. setaceus* throughout the Mexican territory. We used our model to project the scenario in the long term, Figure 3g. iNaturalist data do not record biomass size, so we assigned sizes by random sampling from the Guadalajara data. Our results show that in 200 years from today, 11 % of the Mexican territory will be infested by the weed. This problem could become worse if we consider new introductions due to human activity.

### 2.6 Some methods for weed control

So far, we have analyzed scenarios of weed infestation. At that point, we analyze whether there is some mechanism that can push back or contain the weed. In the first case, we analyzed a territory defined by a 10 × 100 pixel matrix (5 × 50 km). We consider that a border of the territory, defined by a 10 × 10 sub-matrix, is fully infested with a density equal to the median of the Guadalajara data. We track the distance from the infestation *focus* border to the farthest infested pixel over several years. In order to reduce the noise generated by stochasticity, we performed 100 repetitions. As shown in Figure 4a,b, initially the weed grows with velocity of 0.5 km/year, approximately until it reaches a stable velocity of 0.12 km/year, with a mean of 0.13 km/year. Furthermore, to push back the weed, eradication efforts must proceed with a velocity of approximately 0.5 km/year, considering that we know the infested border. However, the cleared territory should be constantly supervised, because the remaining seeds or roots can re-establish the weed in a short time.

**Figure 4.**
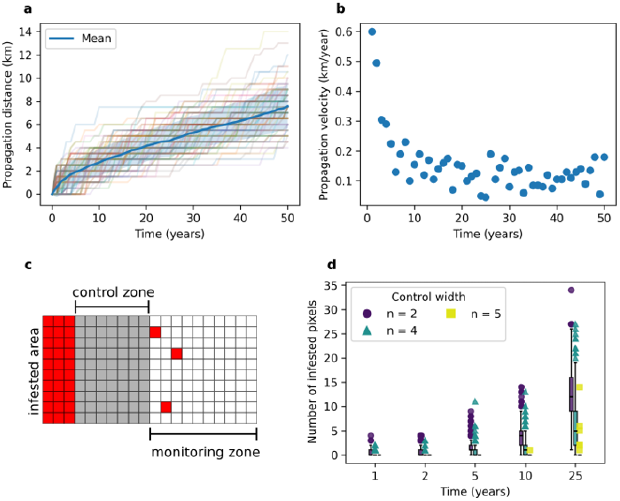
Methods for push back or contain the weed. First we analyze the propagation velocity of the weed over time. In (a), we can observe many simulations (different colors) that, on average, show a smooth growing curve (blue). b. Propagation velocity over time obtained by averaging the velocity of each repetition. We can observe an initial fast propagation than then stabilizes at approximately 0.12 km/year. In order to contains the weed, we analyzed the scenario where we have an initial region (red) infested with *C. setaceus*, a grass-free control area (gray), and a monitoring area (white) in (c). Then, we count the number of infested pixels in the monitoring zone over different time periods (d). The reported number is the average over 1000 realizations. We do not report the case n = 6 because we did not find any infested pixel for this case.

We analyzed the case in which we try to contain the weed by establishing a control stripe of width *n*, with *n* ∈ [2, 4, 5, 6] pixels. We suppose that we can keep the stripe free of the weed (Figure 4c). Then, we calculated the mean number of pixels (over 1000 repetitions) that can cross the control stripe over several time periods (1, 2, 5, 10 and 25 years). As shown in Figure 4c, control stripes of 2 and 4 pixels cannot contain the weed. A 5-pixel-wide stripe contains the weed during the first years, but as time progresses, the probability of crossing increases, and we can observe some infested pixels after 10 or 25 years. Our model predicts that a 6-pixel-wide stripe (3 km) can contain the invasion of the weed. However, we need to take into account that in the control zone we assume that it remains completely free of the weed, over the simulated period. In the field, this is strictly impossible; therefore, perhaps a wider stripe is necessary.

## 3 Discussion

In this work we collected information on the presence of the invasive weed *C. setaceus* in the Mexican territory. From the georeferenced data and biomass size in Guadalajara, we proposed and calibrated a dynamical model of weed invasion. Our model has two principal modules. The first is a seed-dispersal component driven by an exponential spatial probability and the second is a seed-establishing (offspring reaching maturity) module driven by a random local probability given by a threshold. With these two models, we are able to simulate the spatial distribution and biomass size observed at present in Guadalajara, which started in 1996 from planted individuals reaching today the total observed self-propagated ones. However, we observed a contrast between seed viability in the laboratory (87 %) and the establishment of the offspring in the field, reaching maturity (10 %). This is likely due to the several biotic and abiotic factors that determine establishment in the field, considered by our model. So, the urban fabric characterized by asphalt, heavy trampling, municipal weeding and impervious runoff acts as a severe ecological filter that eliminates native competitors, creating highly disturbed niches that accelerate population dynamics and invasive success of *C. setaceus* and, at the same time, limits its huge fertility seed tested in laboratory conditions, maintaining its huge competition against native flora.

Our model predicts the presence of the weed in unsampled points of Guadalajara metropolitan area. We confirmed the accuracy of this predictions, through iNaturalist data, which show records beyond the borders of Guadalajara, just as the model prediction. Once the model was calibrated, we used it to analyze the infestation scenario over medium and long-term periods. Over the medium term, we predict that in 50 years the 1.42 % of the city will be infested. However over 100 years the 51% of the city will be infested according to our model.

iNaturalist data show the presence of the weed throughout the Mexican territory. We used these data to project the scenario over the long term. We found that in 200 years, 11 % of the Mexican territory will be infested. This area is comparable to the current area used for agriculture as well urban and infrastructure lands; actually, the Mexican territory is approximately divided into about 30 % of forest lands, 20 % urban and infrastructure lands and 50 % of agricultural lands (10 % cultivated) [32].

In order to analyze strategies to eradicate or contain the propagation of this weed, we computed the velocity of its invasion. We found that *C. setaceus* could spread at a rate of 0.13 km/year, but at the border of the infested zone, the rate can increase to 0.5 km/year. Then, to eradicate it, there is a need to clear the area at a rate greater than 0.5 km/year. In the field, this represents a formidable challenge because erasing the weed completely, means cleaning the area from seeds and culms with roots. We then analyzed a strategy to contain *C. setaceus* if we assume a completely seed- and culms with roots-free area. We found that a 3-km-wide containment buffer zone is strictly required to halt the invasion front over a 25-year horizon (Figure 4). This operational threshold highlights the critical need for municipal authorities to restrict human activity as an expansion mechanism of the weed, such as the use of *C. setaceus* as an ornamental plant.

Though Mexico formally has a strategy mapped out for the control and eradication of invasive species, the country urgently needs more efforts in scientific research and effective eradication actions [33]. Currently, it is difficult to find municipal regulations governing suitable plant-lists for urban purpose, therefore, even if the plant is cataloged as an high-risk exotic species [34], urbanists can use it in public parks, as a fact happens daily. Similarly, early detection of invasive species in urban environments and timely action would be highly useful to achieve early eradication, thereby avoiding the extremes of Mediterranean European regions already infested by this species [9]. More research is needed on the behavior of this African weed within Mexican territory. Immediate actions must be put into practice to limit its spread—first and foremost by banning its commercialization and propagation in private nurseries, as well as raising awareness among both the civil population and public administrations to halt its cultivation in the country or enforce the use of sterile varieties. The lack of regulated urban vegetation catalogs in fast-growing cities within developing countries turns ornamental green infrastructure into an underrecognized pathway for biological invasion, as exemplified by the unchecked proliferation of invasive species like *C. setaceus*. Otherwise, we will have to familiarize ourselves with this uncomfortable presence in our environment at the expense of native species, just as it has already occurred with the ruby grass [35] in other parts of the world and Northamerica, mainly in the USA and Mexico.

## 4 Methodology

### 4.1 Collected Population Data

The incidence data for *C. setaceus* were collected from Guadalajara, Jalisco, Mexico in 2022, using a random transect survey. We classified the weed into two categories: spontaneous (hereafter: self-

propagated) and human-planted (hereafter: planted). The self-propagated individuals were identified by their random growth in asphalt cracks and on sidewalks (Figure 1 d). The planted individuals were identified by their systematically placement in structured and arranged planters around the urban infrastructure, clearly planted by humans (Figure 1 e). We recorded the number of individuals and the average number of spikes per individual. Due to the growth habit of this species, each bunch of grass measuring approximately 20 to 50 cm diameter was considered as a single individual, to estimate the total invasion in terms of the number of individuals. The coordinates were recorded using a high-precision GNSS mobile tracking with a spatial resolution of approximately 2 meters.

We used the Fitter function from the fitter Python package to find the best statistical distribution for our data. We tested five representative distributions: normal, log-normal, gamma, exponential and beta. We analyzed three cases: total compound data (the two types combined) and separately data by its quality (planted and self-propagated).

Data were rasterized into pixels with a size of 0.5 km. When we binarized the data, three properties emerged: patches, patch size and distances between patches. Together with pixel intensity, we used these properties as a measure of similarity between the real data and the the ones roduced by the mathematical model.

We assumed 1996 as the starting time for the invasion. Therefore, the current status of *C. setaceus* invasion in metropolitan area of Guadalajara (data collected in 2022 [5]) is the result of approximately 26 years of spread.

We also collected records at the present moment in the cities of San Cristóbal de las Casas, Tapachula and Tuxtla Gutiérrez, Chiapas, Mexico, representing an emerging potential invasion. In these last surveys, only the geographic coordinates of each occurrence were recorded; neither the number of individuals nor the number of spikes were recorded, and no distinction was made between planted and self-propagated individuals. Since the observed abundance was considerably lower in Chiapas cities, these data were collected to provide a projection of the future potential spatial pattern of invasion.

To develop a projection of the distribution of *C. setaceus* along the entire Mexican border, we used the iNaturalist dataset of 2,152 total points of interest [20], after our own taxonomic validation based on platform data-points pictures was performed.

### 4.2 Seed productivity, fertility and germination rate

The mature spikes of the species under study were collected from planted individuals (11 spikes in total) and self-propagated individuals (11 spikes in total), from the Guadalajara metropolitan area during both dry and rainy season (as well as February – May and July – October). We counted the spikelets with relative seed number, calculating the mean and standard deviation (Figure 1 f). The average seed number was calculated as the average number of spikes per individual, as recorded during the field trip. Seed fertility was determined using the tetrazolium staining method [36] with 3 replicates of 60 seeds each, under controlled conditions at 24 °C for 48 hours in the dark (Figure 1 g). To determine the germination rate, the seeds were placed in Petri dishes with moist filter paper under the same environmental conditions for 48 hours (Figure 1 h). Three replicates of 60 seeds each were also used.

### 4.3 Mathematical model

Let 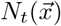 denote the number of spikes of *C. setaceus* at spatial position 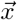 and time *t*. The temporal dynamics are described by a stochastic model based on Monte Carlo simulations. The colonization process is divided into two stages: dispersal and establishment.

Dispersal is modeled by an exponential kernel given by

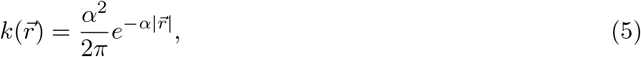

where 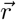 is the displacement vector from the source to the arrival position, and *α* is the dispersal rate parameter. The seed contribution from neighboring sources is given by

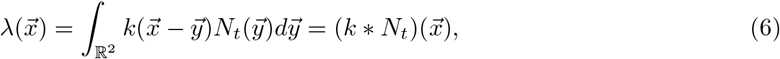

where the symbol * denotes the convolution operator. Thus, considering seed arrivals as a Poisson process with mean 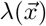, the probability that seeds reach the position 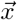 is given by

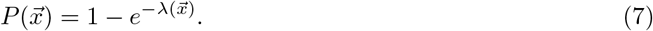

A site is considered a candidate for colonization if 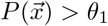, where *θ*_1_ represents an arrival probability threshold.

The seed arrival does not guarantee a successful establishment, it means germination and a potential adult plant. We model establishment by the following mechanism: for each candidate site, a random number *u* ~ *U*(0, 1) is generated and establishment occurs if *u < θ*_2_. So, the parameter *θ*_2_ accounts for biological processes such as germination, seedling survival, and other environmental factors that influence successful establishment.

Once a site is colonized, the initial number of spikes is sampled from a probability distribution *B*, estimated from field observations of naturally established populations of *C. setaceus*. Since local population growth is limited by the carrying capacity, the number of new spikes added at a colonized site is given by

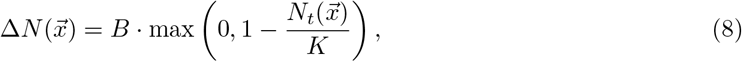

where *K* denotes the carrying capacity. The carrying capacity of the self-propagated population was approximated by the maximum number of planted individuals observed during the field surveys. This assumption is based on the premise that planted individuals are established under favorable environmental conditions and therefore represent the maximum population size that self-propagated individuals could potentially attain.

The biomass is then updated according to

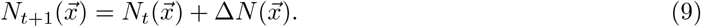

### 4.4 Numerical implementation

We implemented the model on a grid with pixel size of 0.5 km. The temporal step was defined as 1 year. The convolution is computed using the Fast Fourier Transform.

We implemented the simulations in Python. We developed the Python package cenchrus which contains all the functions required to run the simulations [37].

### 4.5 Model optimization

In order to accurately reproduce the current state of *C. setaceus* in the Guadalajara metropolitan area, we estimated the thresholds *θ*_1_, *θ*_2_ and the probability distribution parameters by comparing the output of our model after 26 years time steps, using the rasterized planted matrix as the initial condition, with the rasterized self-propagated matrix. For both observed and simulated pattern, we compute the vector of summary statistics:

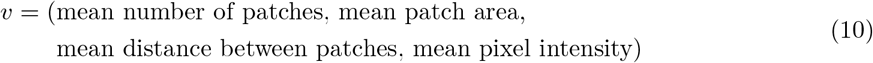

and defined the objective function as

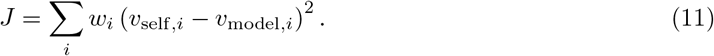

where *v*_self,*i*_ and *v*_model,*i*_ denote the *i*th components of the summary statistic vectors for the self-propagated and simulated pattern, respectively. The coefficients *w*_*i*_ are the weights assigned to each summary statistic according to its relative importance in the optimization. The weights were selected empirically through a trial-and-error procedure. Preliminary experiments showed that the model readily reproduced the mean pixel intensity and the mean distance between patches, while reproducing the mean patch area was more challenging, and accurately matching the number of patches proved to be the most difficult. Consequently, larger weights were assigned to the latter quantities. The final weights used in the optimization were *w* = (0.8, 0.15, 0.025, 0.025).

To reduce the variability introduced by the Monte Carlo simulation, the summary statistic vector was averaged over 100 independent realizations at each optimization step. These realizations were evaluated in parallel to improve computational efficiency. Furthermore, because the presence records were not exhaustive, *i*.*e*., many pixels recorded as zero do not necessarily indicate the absence of the species but rather unexplored areas, the objective function was computed only for pixels within the concave hull (ratio = 0.1), which was used to tightly enclose the surveyed area.

We implemented optimization with the minimize function of scipy.optimize Python package via the Nelder-Mead method. Geographic figures were obtained using Python-based processing and/or QGis 4.2.2 with the Mexican administration layers of INEGI [38]. These geographic data are publicly available and freely downloadable (associated github repository).

## Acknowledgments

R. R. thanks Secretaría de Eduación, Ciencia, Tecnología e Innovación de la CDMX (SECTEI) for project 5613c25.

